# Continuous automated analysis workflow for MRS studies

**DOI:** 10.1101/2022.11.03.515056

**Authors:** Helge J. Zöllner, Christopher W. Davies-Jenkins, Erik G. Lee, Timothy J. Hendrickson, William T. Clarke, Richard A. E. Edden, Jessica L. Wisnowski, Aaron T. Gudmundson, Georg Oeltzschner

## Abstract

**Background:** Magnetic resonance spectroscopy (MRS) can non-invasively measure levels of endogenous metabolites in living tissue and is of great interest to neuroscience and clinical research. To this day, MRS data analysis workflows differ substantially between groups, frequently requiring many manual steps to be performed on individual datasets, e.g., data renaming/sorting, manual execution of analysis scripts, and manual assessment of success/failure. Manual analysis practices are a substantial barrier to wider uptake of MRS. They also increase the likelihood of human error and prevent deployment of MRS at large scale. Here, we demonstrate an end-to-end workflow for fully automated data uptake, processing, and quality review.

**New Method:** The proposed continuous automated MRS analysis workflow integrates several recent innovations in MRS data and file storage conventions. They are efficiently deployed by a directory monitoring service that automatically triggers the following steps upon arrival of a new raw MRS dataset in a project folder: (1) conversion from proprietary manufacturer file formats into the universal format NIfTI-MRS; (2) consistent file system organization according to the data accumulation logic standard BIDS-MRS; (3) executing a command-line executable of our open-source end-to-end analysis software Osprey; (4) e-mail delivery of a quality control summary report for all analysis steps.

**Results:** The automated architecture successfully completed for a demonstration dataset. The only manual step required was to copy a raw data folder into a monitored directory.

**Comparison with Existing Method(s):** The workflow presented here is the first implementation of a continuous automated MRS analysis ecosystem based on NIfTI-MRS and BIDS-MRS standards. Traditional MRS workflows are non-standardized, often require manual input, and frequently noy compatible with established neuroimaging workflows. This workflow should therefore facilitate integration of MRS into large-scale and multi-center studies.

**Conclusions:** Continuous automated analysis of MRS data can reduce the burden of manual data analysis and quality control, particularly for non-expert users and multi-center or large-scale studies.

## Introduction

Magnetic resonance spectroscopy (MRS) provides access to measuring levels of endogenous biochemicals in vivo with a clinical MRI scanner. Approximately 15 different metabolites can be resolved, e.g., markers of neuronal integrity and cell proliferation, neurotransmitters, and antioxidants. MRS is therefore a method of great interest to clinical research and neuroscience. However, although MRS predates other modern MRI modalities (fMRI, DTI) by a decade, it is not used as widely (1). MRS analysis workflows are commonly not automated and require local spectroscopy expertise. Everyday use of MRS has therefore remained restricted to a few specialized research centers.

One barrier to efficient inclusion of MRS into multimodal imaging protocols is the number of manual steps required. Applied research studies with MRS in the acquisition protocol require manual data renaming and sorting, manual amendment of analysis scripts, manual execution of the analysis pipeline, and manual assessment of processing success on individual datasets. Depending on the amount of data and complexity of the protocol, these tasks can take a substantial amount of time. Manual analysis labor is therefore a strongly rate-limiting factor that prevents the deployment of MRS at scale. Finally, the requirement for manual data analysis increases the likelihood of human error, e.g., copying the wrong files, using incorrect filenames, or executing the wrong analysis script.

Because MRS data are less familiar than MRI (and because spectra are not brain images), it can be challenging for non-expert users to distinguish between ‘successful’ and ‘unsuccessful’ acquisitions at the scanner. It is also regrettably common for data to be corrupted or even lost entirely due to procedural error (e.g., a technologist uses an incorrect protocol version, changes a sequence parameter, or accidentally places an MRS voxel in the wrong hemisphere) or as a consequence of hardware and software updates (e.g., a software update introduces a new default exam parameter that overrides an acquisition-critical one). Regular data inspection is therefore necessary to capture sudden unintended acquisition protocol changes – while our recommendation to users is “check after every scan”, this places a large burden on research staff, with the result that it is not consistently followed.

In this manuscript, we describe a fully automated data review and end-to-end analysis workflow to support non-expert users of MRS conducting local research studies. This procedure makes use of several recent innovations in MRS file formats and large-scale neuroimaging data management, which make it much easier to deploy repeated analyses. The workflow can be customized to accommodate local project protocols and IT infrastructure.

## Methods

The primary goal of this work is to reduce the manual overhead involved in checking data regularly, and to make modern MRS analysis easier. The secondary goal is to demonstrate the benefits of automated analysis workflows for the deployment of MRS methods at scale. Reducing manual analysis effort is necessary to open MRS methods up to be a part of multi-site consortium projects or continuous long-time local initiatives.

First, a command-line job scheduling or file-watching utility is deployed at regular intervals to monitor the content of the project folder. If the service detects previously unencountered raw MRS data, a script is launched to convert them into a standardized folder and file structure according to a pre-defined study-specific template. Next, a new analysis job is automatically set up and executed. After completion of the analysis, the tool sends the user an e-mail with a report summarizing quantitative results, quality control criteria, and analysis diagnostics (warnings, errors). This effectively removes the need to check data manually after every scan as the workflow generates the email report for every new dataset automatically – if no email is received even though an MR scan was performed, the user is alerted to check whether data export from the scanner occurred as intended. Equally, if the data exported are incomplete (i.e., do not agree with the expected data specified in the study template), the email report will signal the need to repeat data export.

The workflow (**Figure 1**) integrates several common processing steps with MRS-specific data organization and workflow innovations:

scripted raw data format conversion from proprietary manufacturer file formats into the universal storage format NIfTI-MRS
consistent organization and metadata annotation according to the data accumulation logic standard BIDS
a compiled command-line executable version of our open-source end-to-end analysis software Osprey
regular monitoring of a raw data project folder for automation of the above steps when a new dataset arrives
e-mail notification upon completion of a new automated analysis.

**Figure 1:**
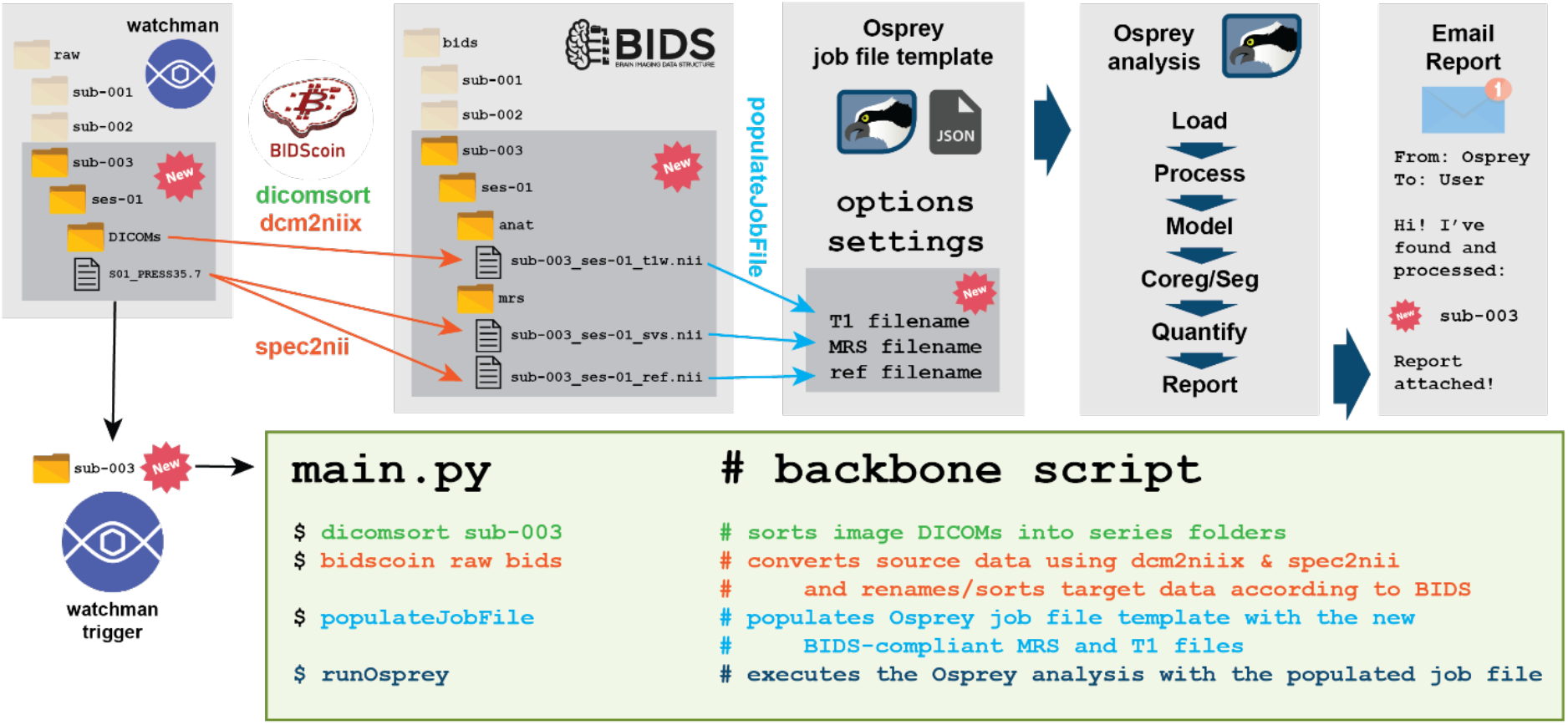
Illustration of the automated MRS data analysis workflow. The *watchman* service notices a new folder in the raw data directory that it monitors. It triggers the backbone script, which first uses *dicomsort* to sort image DICOM into series folders, then executes *bidscoin* which uses the *dcm2niix* and *spec2nii* programs to convert the raw data to NIfTI/NIfTI-MRS/BIDS format. The script then populates a template job file with paths to the new data and passes the job file on to an *Osprey* analysis instance. The last *Osprey* step sends a confirmation e-mail to specified addresses.

### Source data directory monitoring

The foundation of the automated workflow is the source data directory monitoring, which was implemented using an open-source file watching service (*watchman*) to continuously monitor the source data directory for changes. When *watchman* detects an added subject-level folder in the source data directory, it performs a quick check whether this new folder fulfills source data formatting requirements. If it does, watchman triggers the backbone script *main.py*.

### Backbone script

The ‘chain of commands’ (raw data conversion, analysis execution, reporting) is strung together in a small Python script (*main.py*). It requires no modification by the user and works for all raw data formats and sequence types supported by *spec2nii* and *Osprey*. In addition to the automated directory monitoring, the *main.py* script can also be triggered manually. Each step of the workflow integrated in the *main.py* script is described in detail below.

### NIfTI-MRS & BIDS-MRS conversion

This standardized MRS file format convention was developed to overcome longstanding issues with different raw data formats (2). Each manufacturer uses several proprietary file types (Siemens DAT and RDA; Philips SDAT/SPAR, DATA/LIST and SIN/LAB/RAW; GE .7 P-file; multiple vendor flavors of DICOM). These contain different non-standardized and mutually incompatible data and header information and require separate code to parse. NIfTI-MRS is based on the NIfTI-2 standard that is widely used in neuroimaging. Raw data is stored in well-defined array dimensions and accompanied by standardized header information fields saving all metadata necessary to reconstruct the spectra. The Brain Imaging Data Structure (BIDS) is a neuroimaging data organization standard (3). File and folder naming conventions create a standardized hierarchy (study, subject, session, modality) that benefits the automation of analysis pipelines and data sharing. BIDS further catalogues required metadata on technical details and demographics. BIDS-MRS is an MRS-specific extension of BIDS currently in development (4).

Several tools have been developed to automate the conversion and sorting of raw DICOM image source data into BIDS-compliant NIfTI, e.g., HeuDiConv (5), dcm2bids (6), bidskit (7), etc. They use heuristics based on DICOM metadata and file paths to recognize image types and determine ‘maps’, i.e., well-defined procedures for BIDS-compliant file naming and metadata annotation. While very flexible and powerful, their learning curves can be steep. We decided to use BIDScoin (8) for two reasons:

1. its user-friendly GUI allows researchers to define appropriate mapping strategies after the software has made intelligent guesses.
2. it includes a plugin to run spec2nii, a command-line conversion tool from vendor-proprietary formats into NIfTI-MRS.

The backbone script first calls *dicomsort*, a command-line utility included in BIDScoin that sorts unsorted image DICOMs into folders named for the image series they were acquired under, e.g., “T1-MPRAGE”. The BIDScoin GUI (*bidsmapper raw bids*) is used to create a bidsmap file to automatically recognize the source MRS dataset and the source anatomical image DICOMs in these image series folders. This initial mapping can, for example, be carried out on a pilot dataset of a specific study. Using *bidseditor raw*, the user defines output naming heuristics, including identifying the DICOMs as a, for example, T1-weighted image, and adding BIDS-MRS naming labels and JSON header entries to identify the voxel location, localization techniques and MRS sequence types. These translation heuristics, from source to BIDS-compliant output, are stored in a YAML-formatted file (the *bidsmap*) that serves as a template for any subsequently acquired subject folders (*sub-002, sub-003*, etc.). This template is generated once for each study.

### Osprey

Osprey is our free modular open-source software written in MATLAB that completes all consensus-recommended modern MRS data analysis steps, i.e., pre-processing of raw data, modeling pre-processed data with linear-combination modeling, and converting model parameters into metabolite concentration measures (9). Osprey performs comparable to similar linear-combination analysis tools (10,11). Osprey can analyze conventional and edited (single-or multi-metabolite) MRS data and includes simulated basis sets for all major vendors, localization techniques, and metabolites. Custom basis sets that were simulated with external tools (12–14) can easily be integrated.

For the present work, we compiled Osprey v2.4.0 into standalone executables for Windows 10 and MacOS using the MATLAB Compiler. We further created an installation routine that automatically downloads and installs the necessary MATLAB Runtime libraries. The compilation script is freely available and allows users to generate executables for other operating systems.

Osprey requires a JSON-formatted ‘job file’ specifying the analysis options, settings, and the complete paths to the NIfTI-MRS/BIDS-formatted data. This workflow automatically generates this file by populating a template job file with the newly encountered BIDS-formatted data. Analysis options and settings are controlled by a *master_settings.json* file which is created once for each study. Once the job file has been automatically generated, the workflow calls the Osprey executable, with the job file as the only input argument.

### Reporting

Once Osprey has completed the requested analysis, a standardized reporting page (in HTML format), which is part of the standard Osprey analysis, is packaged into a zip file and attached to an e-mail sent to a set of addresses specified by the user in the job file. The report contains visualizations of the different analysis stages (raw data loading, pre-processing, modeling, co-registration, segmentation) and quality control metrics (FWHM, SNR, fit error, frequency drift) as described in a recent consensus paper (15). The information in this report is a condensed single page summary of the extensive visualization available in the Osprey GUI and the routine Osprey PDF output. The report page does not contain protected health information (PHI, as defined by the U.S. Health Insurance Portability and Accountability Act, HIPAA) and can therefore be shared directly. Further, the HTML format allows for direct integration into webbased data management systems (16), and is therefore particularly suitable for multi-modal multi-center studies (MRS, MR imaging, clinical, behavior, genetics, etc.).

### Availability of the workflow

The implemented workflow including a documentation, the example data and scripts used in the manuscript, compiled Osprey, and all additionally needed code is freely available in a GitHub repository (https://github.com/HJZollner/ContinuousAnalysisMRS).

### Deployment requirements

The Python-based BIDScoin application and the Osprey MATLAB executables can be installed on virtually any platform. This includes local workstations, institutional workstations or cloud-based servers. The *watchman* service for file watching is freely available for UNIX-type systems, but could be replaced with system-specific scheduling systems such as *cron* or *Windows Task Scheduler*. Finally, a local workflow to transfer the raw data from the scanner into the monitored raw data folder has to be established.

### Local test implementation and test dataset

We installed the fully automated workflow on a Windows workstation (Windows 11). The raw data and BIDS output folders were created on a cloud-based storage system (OneDrive Business).

Next, we tested the workflow on raw data from a single site with a 3T General Electric scanner from the publicly available Big GABA dataset (site ID ‘G1’) including a total of seven subjects (17–19). Specifically, the image DICOMs of a 3D anatomical T1-weighted MP-RAGE scan and a short-TE (TE = 35 ms) PRESS scan in .7 format (20) were processed.

Using the BIDScoin GUI and an example dataset (*sub-001*), we created a bidsmap file to automatically recognize the source MRS dataset and the source anatomical image DICOMs in these image series folders. Using *bidseditor raw*, we defined output naming heuristics, including identifying the DICOMs as a structural T1-weighted image, and adding BIDS-MRS labels *vox-pcc* to denote the voxel location in the posterior cingulate and *acq-press* to highlight that these data were acquired with PRESS localization. These translation heuristics served as a template for the remaining subjects.

Finally, the analysis options and settings in the master_settings.json file were updated according to the prerequisites of the test dataset (short-TE data, no eddy-current correction, and local email address). Afterwards, each subject’s data were transferred into the monitored raw data folder to mimic ongoing data acquisition to test the automated workflow.

## Results

The workflow presented here was successfully deployed to process the test dataset. After the initial configuration of the bidsmap, Osprey job file templates, and *watchman* scripts, no further user intervention was required. All seven single-subjects datasets were automatically pushed through the entire pipeline (DICOM sorting, NIfTI-MRS/BIDS conversion, Osprey analysis). Osprey output files included the summary report sent via e-mail (**Figure 2**). A full example report can be found in **Supplementary Material 1** (converted to PDF format to comply with journal guidelines). The routine file system output generated by Osprey (PDF visualizations, tab-separated values files holding metabolite estimates for further analysis, binary voxel masks, etc.) is automatically saved into a *derivatives* folder as specified by the parent BIDS. More specifically, this *derivatives* subdirectory is created in the root directory of the source BIDS dataset.

**Figure 2:**
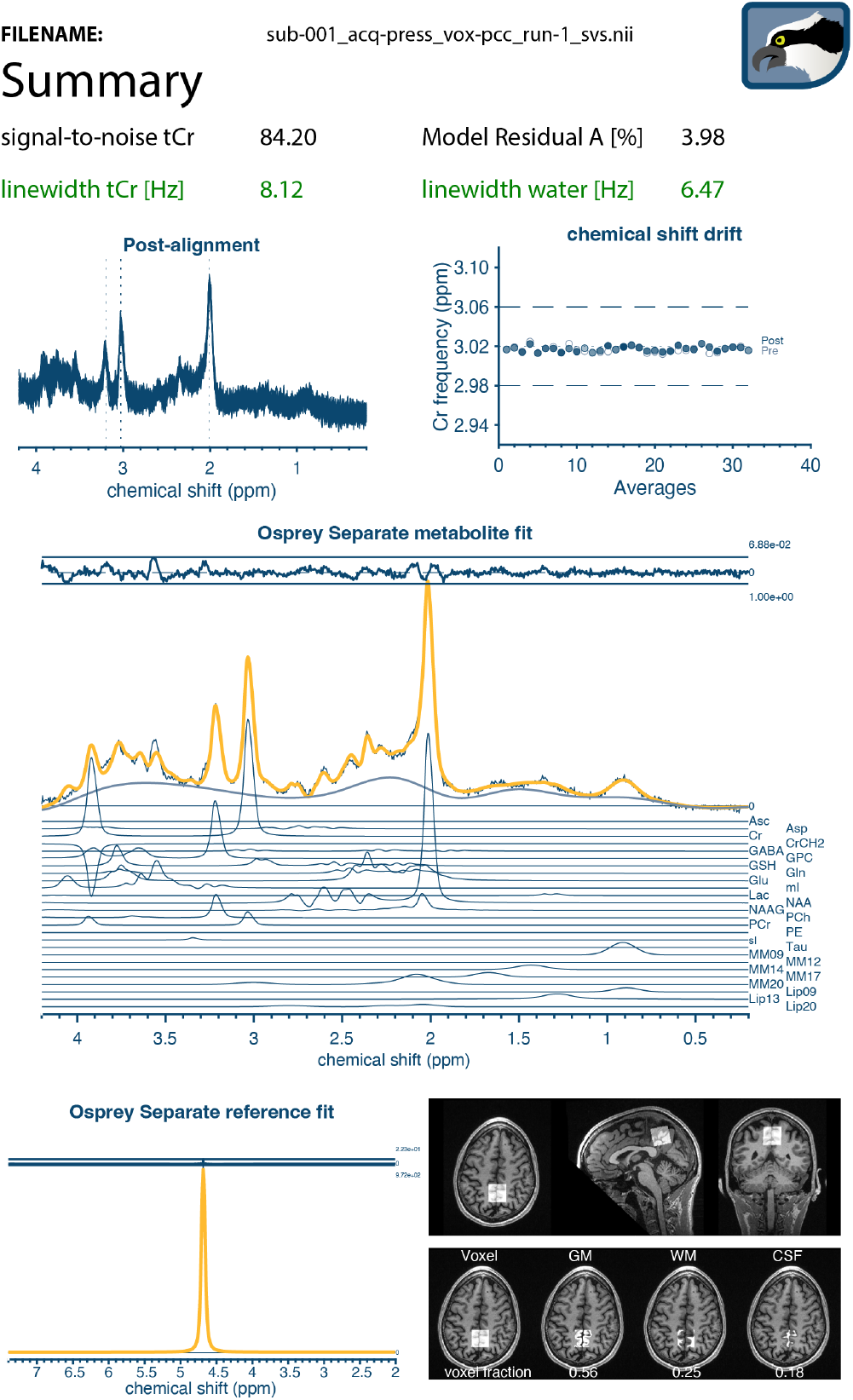
Summary report on all key steps of the automated MRS data analysis workflow (lightly edited from HIPAA-compliant HTML output to minimize whitespace for presentation purposes). The HTML report is automatically e-mailed to a user-specified address after completion of the Osprey analysis. The summary includes quantitative quality control metrics (linewidth, SNR, model fit error) and visualizations of the processing, modeling, and co-registration/segmentation steps of the automated Osprey pipeline.

## Discussion

We have demonstrated an example of an automated workflow for project folder organization, data analysis and reporting for clinical MRS studies. Automation of MRS data analysis can improve reproducibility and efficiency, particularly at sites that historically have not established local workflows. The immediate feedback helps users identify sudden disruptive changes to the hardware or software setup, for example after scanner upgrades. Lastly, automated data analysis can lower the entry-level threshold to engage with MRS in general, and provide much-needed scale.

Standardized workflows and data storage conventions also increase reusability of existing analysis code at various levels. Once set up for a particular study, the setup described herein can easily be ported to a new study protocol, simply by adjusting the bidsmaps and Osprey job file templates. This increases ‘horizontal’ efficiency for the individual researcher, between members of a research group, and between locations of multi-site consortia, but also ‘longitudinal’ efficiency for incoming personnel taking over responsibility for existing projects.

This automated analysis workflow may also serve as a template for the integration of MRS into large-scale multi-center multi-modal neuroimaging studies. MRS data acquisition and analysis have not been easy scalable in the past since they were limited by the amount of manual effort they demand. As a result, even the largest MRS datasets may only include a few hundred subjects – orders of magnitude less than the most powerful databases of structural, functional, or diffusion imaging. BIDS has transformed the way in which large-scale neuroimaging datasets are made accessible to other researchers, and the anticipated publication of the BIDS-MRS extension will open up many existing neuroimaging repositories for use with MRS data.

### Alternative strategies

Neuroimaging centers may already have solutions to transform source imaging data into BIDS format in place, e.g., for fMRI, PET, or EEG data. The BIDScoin part of the workflow presented here may easily be swapped for existing ‘BIDS-ification’ methods, although it offers the advantage of already offering a work-in-progress MRS integration via a *spec2nii* plugin.

We demonstrate the workflow with Osprey, our own software for MRS pre-processing, modeling, and quantification. Researchers can, of course, swap Osprey for any other MRS analysis software that allows command line execution. Note, however, that full BIDS compliance requires NIfTI-MRS as the core file storage format for data input and output. NIfTI-MRS is supported by recently published open-source software packages (FSL-MRS (21), spant (22)) and the latest versions of some seasoned ones (Vespa (23), jMRUI (24)). LCModel (25), on the contrary, is unlikely to directly interface with NIfTI-MRS unless data are prepared with modular processing packages like FID-A (26) which includes NIfTI-MRS reading and writing routines. Osprey also includes a fully-functioning wrapper to call LCModel. Such modifications can be achieved by simply modifying the *main.py* backbone script to integrate other analysis software. Additionally, any summary reports have to be integrated into the current zip file to be available in the email alert. Researchers wishing to maintain vendor-native formats (SDAT/SPAR, RDA, etc.) may construct similar workflows involving directory monitoring and standardized file naming conventions, although they forfeit many of the advantages that BIDS offers through highly formalized storage logic and existing software solutions.

Finally, continuous directory monitoring is an optional part of the workflow. We selected the *watchman* service because it is free, open-source, and compatible with different OS architectures. Researchers may opt for similar file monitoring services (*fswatch*), scheduled execution (*cron* on Unix-type systems, *Task Scheduler* on Windows) at larger time intervals, or wish to execute the workflow manually.

### Limitations and future perspectives

Integration of MRS into BIDS and its software ecosystem is a recent innovation that is continuously evolving. NIfTI-MRS conversion with *spec2nii* is supported for many common formats and sequence implementations, but may require further modification for custom sequences, either directly in the source code or with command line options to specify data dimensions and add metadata. Similarly, BIDScoin currently only supports a single ‘type’ of MRS data per bidsmap, i.e., two different bidsmaps need to be defined to accommodate study protocols with, e.g., one PRESS and one MEGA-PRESS dataset. In this case, the backbone script *main.py* needs to feature two instances of *bidscoiner* with the respective bidsmaps as input argument.

The filename mapping heuristics can be tailored to be powerful and versatile thanks to regular expressions in the filters, but still require a relatively consistent study protocol to be executed on the scanner. If sequence parameters, scan sequence names and order, and other data attributes and properties change frequently, the BIDScoin heuristics are likely to fail. This workflow is therefore primarily useful for long-running application studies with a fixed protocol, but less suitable for exploratory methodological piloting, which typically requires custom manual analysis anyway.

Quality control metrics remain a challenge for MRS. Model residuals, Cramer-Rao Lower Bounds, spectral linewidth, and signal-to-noise ratio all provide valuable information, but they often fail to reliably indicate the presence of major artefacts like lipid contamination or out-of-voxel echoes. Visual inspection of individual spectra therefore continues to be a key benchmark by which data quality is judged. This practice is clearly not scalable for large-scale studies or even single-site spectroscopic imaging. CSI/MRSI produces thousands of spectra per dataset, and the output is still often judged by the visual appearance of the final color map. Several quantitative metrics for quality assessment have been proposed (27–30), but have not been tested or implemented in mainstream MRS analysis. Automated scalable workflows like the one described here may facilitate uptake and benchmarking of these quantitative QC metrics. After establishing QC metrics, a QA pass/fail indicator could be integrated into the header of the report and the overarching database that gather the study results, such as LORIS (16).

This manuscript is focused on the general presentation of the automated workflow and was only tested for a single vendor and MRS sequence type, but should generalize well to other vendors and acquisitions. *Bidscoin* and *spec2nii* generate a vendor-independent standardized file structure and MRS data format. The current spec2nii implementation already supports MRS data from all major vendors and various multi-dimensional MRS experiments. Higher dimensions may include coils (for raw uncombined data), dynamics/repeats (temporal transients), and additional encoding techniques (edited, 2D, diffusion-weighted MRS, etc.). Similarly, Osprey’s flexible implementation allows for MRS analysis of conventional, metabolite-nulled, and edited MRS. It has been benchmarked against other algorithms and is tested extensively with multi-site and multi-vendor data. Additionally, all three tools are actively developed, public, open-source resources and therefore easily adapted for potential novel MRS techniques in the future.

## Conclusion

In summary, we present an open-source end-to-end workflow for continuous automated analysis of MRS data, making use of recent advances in standardized file format and data storage conventions (NIfTI-MRS, BIDS-MRS). Reduced manual analysis overhead can help simplify the integration of MRS into large-scale multi-modal imaging studies and clinical trials.

## CRediT author statement

**Helge J. Zöllner**: Formal Analysis, Writing – Review & Editing, Visualization. **Christopher W. Davies-Jenkins**: Writing – Review & Editing. **Erik G. Lee**: Software Implementation, Writing – Review & Editing. **Timothy J. Hendrickson**: Software Implementation, Writing – Review & Editing. **William T. Clarke**: Software Implementation, Writing – Review & Editing. **Richard A. E. Edden**: Writing – Review & Editing, Project administration, Supervision, Funding acquisition. **Jessica L. Wisnowski**: Writing – Review & Editing, Supervision. **Aaron T. Gudmundson**: Software Implementation, Writing – Review & Editing. **Georg Oeltzschner**: Conceptualization, Methodology, Writing – Original Draft, Writing – Review & Editing, Project administration, Supervision, Funding acquisition.

## Acknowledgements

This work has been supported by NIH grants R00 AG062230, R21 EB033516, R01 EB016089, R01 EB023963, and P41 EB031771. WTC is supported by funding from the Wellcome Trust [225924/Z/22/Z]. JLW is supported by NIH grants U01 DA055362 and K23 HD099309. The authors would like to thank Dr. Marcel P. Zwiers (Centre for Cognitive Neuroimaging, Donders Institute for Brain, Cognitive and Behaviour, Radboud University) for his constructive feedback on the implementation of MRS in *bidscoin*.

