## Supplementary Material 1 for "Continuous automated analysis workflow for MRS studies"

Osprey Analysis Report

**DATE:** 01-Feb-2023 **FILENAME:** sub-001_acq-press_vox-pcc_run-1_svs.nii

### Summary:

**signal-to-noise tCr** 84.20

**Model Residual A [%]** 4.05

**linewidth tCr [Hz]** 8.12

**linewidth water [Hz]** 6.47

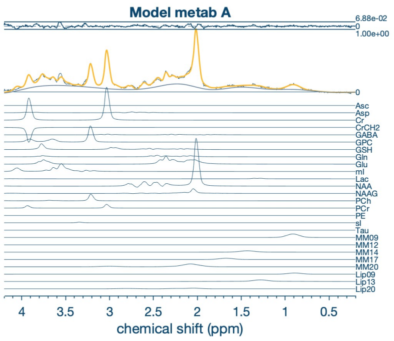


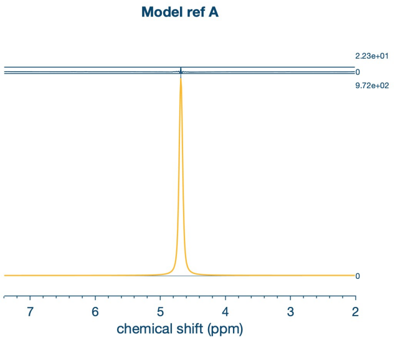


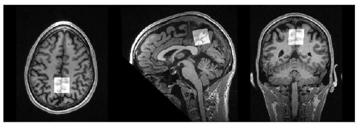

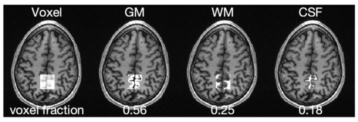


### Results of spectral registration:

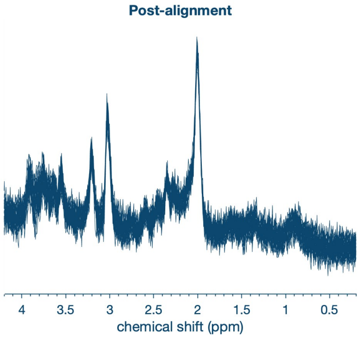


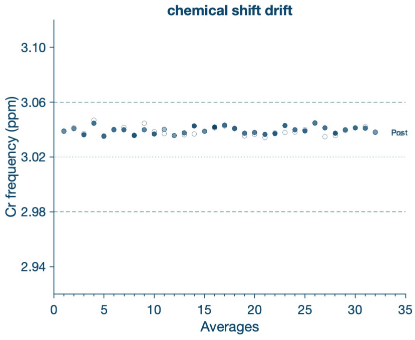


### Averaged spectra:

**signal-to-noise tCr** 84.20

**linewidth tCr [Hz]** 8.12

**linewidth water [Hz]** 6.47

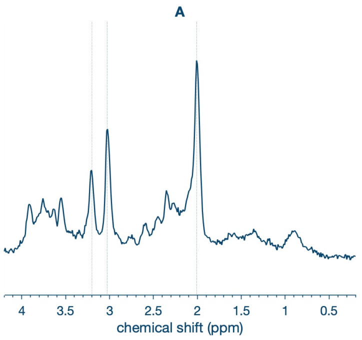


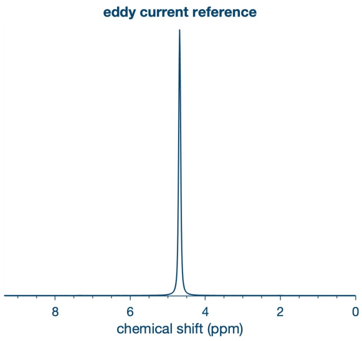


### Model results:

**Relative Model Residual A [%]** 4.05

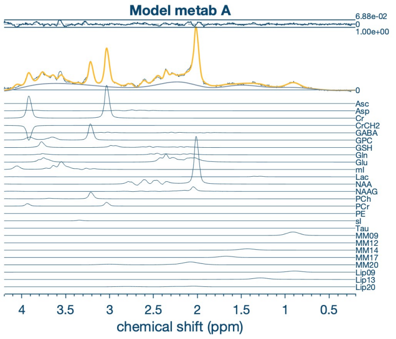


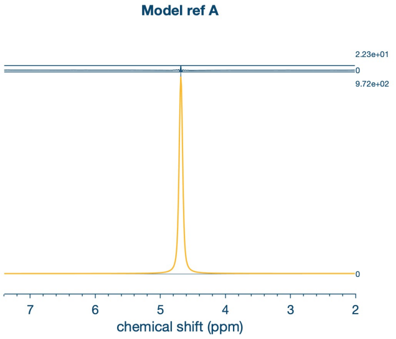


### Coregistration & Segmentation:

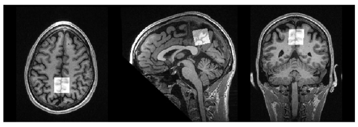

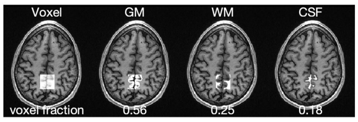


Supplementary Material 1 - HIPAA-compliant HTML Summary report on all key steps of the automated MRS data analysis workflow . The HTML report is automatically e-mailed to a user-specified address after completion of the Osprey analysis. The summary includes quantitative quality control metrics (linewidth, SNR, model fit error) and visualizations of the processing, modeling, and co-registration/segmentation steps of the automated Osprey pipeline.
